# Amplified DNA Heterogeneity Assessment with Oxford Nanopore Sequencing Applied to Cell Free Expression Templates

**DOI:** 10.1101/2024.06.02.597048

**Authors:** Sepehr Hejazi, Afrin Ahsan, Mohammad Kashani, Nigel F Reuel

## Abstract

In this work, Oxford Nanopore sequencing is tested as an accessible method for quantifying heterogeneity of amplified DNA. This method enables rapid quantification of deletions, insertions, and substitutions, the probability of each mutation error, and their locations in the replicated sequences. Amplification techniques tested were conventional polymerase chain reaction (PCR) with varying levels of polymerase fidelity (OneTaq, Phusion, and Q5) as well as rolling circle amplification (RCA) with Phi29 polymerase. Plasmid amplification using bacteria was also assessed. By analyzing the distribution of errors in a large set of sequences for each sample, we examined the heterogeneity and mode of errors in each sample. This analysis revealed that Q5 and Phusion polymerases exhibited the lowest error rates observed in the amplified DNA. As a secondary validation, we analyzed the emission spectra of sfGFP fluorescent proteins synthesized with amplified DNA using cell free expression. Error-prone polymerase chain reactions confirmed the dependency of reporter protein emission spectra peak broadness to DNA error rates. The presented nanopore sequencing methods serve as a roadmap to quantify the accuracy of other gene amplification techniques, as they are discovered, enabling more homogenous cell-free expression of desired proteins.

## Introduction

Cell-free expression (CFE) has become an established technique for rapid protein prototyping and relies on pure DNA inputs to achieve maximal expression of a protein of interest[1]. Titers from CFE reactions have improved significantly to make this method capable of screening and manufacturing custom proteins for sensors, diagnostics, therapeutics, functional materials, and biocatalysts [2, 3]. CFE reactions require cell extract, supplements, and a genetic template to perform transcription, translation, and some post-translational modification. The homogeneity of the final protein depends highly on the quality of the genetic template [4]. The most common template is circular dsDNA (plasmid) due to its stability and compatibility with cell-based amplification and expression techniques. However, CFE can also use linear expression template (LET) genes. These genes can be sourced from chemically synthesized DNA, followed by a conventional polymerase chain reaction (PCR) amplification [5]. Although protein production levels from LET are generally lower than plasmids, they do enable rapid protein prototyping for protein discovery as it can be completed on a time scale of hours rather than traditional plasmid preparation which takes days to clone, transfect, and amplify [6].

LET amplification quality is governed mainly by the proof-reading quality of the polymerase used. Many polymerases possess 3’-5’ exonuclease activity which is used for proofreading, that is, the removal of incorrectly generated nucleotides after further extension in the exo site and then moving the DNA back to the polymerase site [7]. It is important to note that DNA polymerases have a balance between the speed and accuracy of DNA replication. Excessive proofreading may slow down the rate of replication and increase the risk of DNA damage due to exposure to reactive oxygen species and other mutagens [8]. On the other hand, insufficient proofreading can increase the frequency of mutations and lead to genetic instability [9]. Taq DNA polymerase is a commonly used enzyme for PCR due to its early use in PCR protocols and superior thermal stability than previous DNA polymerases such as DNA Polymerase I [10].

Comparisons of fidelity between polymerases can be quantified in absolute terms, typically by determining the number of errors per 1,000 or 10,000 nucleotides or the probability of error per base pair. Also, fidelity can be expressed in relative terms by using Taq DNA polymerase as the reference standard and comparing other polymerase fidelities to this enzyme [11, 12].

Higher fidelity polymerases have been engineered to improve the error rates caused by PCR amplification, such as Q5 High-Fidelity DNA Polymerase and Phusion High-Fidelity DNA Polymerase which have 280 and 38 times higher fidelity than Taq, respectively[12]. Rolling circle amplification (RCA) is a variation on traditional PCR, in that the linear template is circularized and a strand displacement active polymerase (e.g., Phi29) continuously amplifies the template into a long concatemer. This technique can amplify a start template more than 1000 times (achieving 10-20 μg DNA from 10 ng of initial DNA template) and has been demonstrated to be compatible with CFE [13]. Again, this is done with a level of fidelity, benchmarked at around 5 times higher than Taq DNA polymerase due to the Phi29 DNA polymerase fidelity[14].

To assess the amplification fidelity, gene sequencing technologies can be used to quantify amplification error rates. Traditional Sanger sequencing can be used[15], however, massively parallelized, next generation sequencing (NGS), which is also known as second generation, (e.g., Illumina) provide a larger data set for error analysis. One limitation to this approach is the use of short reads (typically 75-300 bp) which requires a complex genome assembly that works well to reconstruct common sequences (ideally a single sequence) but lacks the depth to capture mutations on pieces that fail to align. To address the limitations of NGS, the emergence of third-generation sequencing technologies, exemplified by PacBio and Oxford Nanopore Technologies (ONT), has provided solutions for high-throughput and also long-read sequencing, with PacBio achieving up to 20 kilobase pairs (Kbp) and ONT reaching up to 1 mega base pair (Mbp) read-length limits. This feature facilitates error assessment at the individual molecule level, making them ideal sequencing methods for this research purpose [16]. While PacBio offers higher read accuracy compared to ONT, the capital cost of the system required for PacBio sequencing poses challenges for typical biotech labs, limiting accessibility and necessitating the shipment of samples for sequencing. In contrast, the ONT MinION device is comparatively more affordable and remarkably compact. It is compatible with various sequencing techniques, including barcoded, rapid, and ultra-long DNA/RNA sequencing [17]. A current limitation for ONT is the higher error rates of nucleotide identification compared to Illumina, however this is partially mitigated by the vast quantity of reads capable from a single sample in Oxford Nanopore sequencing run (10–15 gigabases (Gb) of DNA reads in 48 h on a minion device) [18]; this large amount of independent long sequence reads could enable measurement of population shifts in the number of insertions, deletions, and mutations with statistical certainty. Also, recently a new chemistry kit (LSK114) for ONT was introduced which can sequence a large fragment of DNA input with the read accuracy of more than 99% (30.1% of the reads achieving more than 99% and 98.34% in average) auguring even greater accuracy with future platform improvements[19]. Here we demonstrate heterogeneity analysis of amplified gene templates using Oxford Nanopore sequencing and frame its use with quality assessment of amplified templates used in CFE.

## Materials and Methods

### Gene amplification and characterization workflow

A general overview of the experimental workflow is presented first (**Figure 1**), followed by details on each part. The pJL1-sfGFP Plasmid (Addgene Plasmid #102634) was used as our standard expression template due to its widespread use in the CFE community as a clear functional benchmark – the encoded super folder green fluorescent protein (sfGFP) has robust translational efficiency and can be easily screened by fluorescence output. The plasmid was transfected in *E. coli* TOP10 cells and purified. Linear sfGFP gene fragments were designed from this plasmid, amplifying everything between the T7 promoter and termination regions; additional bases were added to allow for circularization and rolling circle amplification (supporting information section 1). Cleaned, purified, and linearized DNA was repaired and end-prepared (poly-A tailing) for adapter ligation, attaching the motor protein for nanopore sequencing to the DNA fragments. The DNA library was then loaded and sequenced with the Oxford Nanopore minION chip. Output files were base called and aligned with the reference sequence for the sfGFP gene and custom codes were used to extract the mutation data from it. Normalized mutation (substitution, insertion, deletion) error rates were the final output of this workflow (**Figure 1**). Additional details for each of these steps follow.

**Figure 1.**
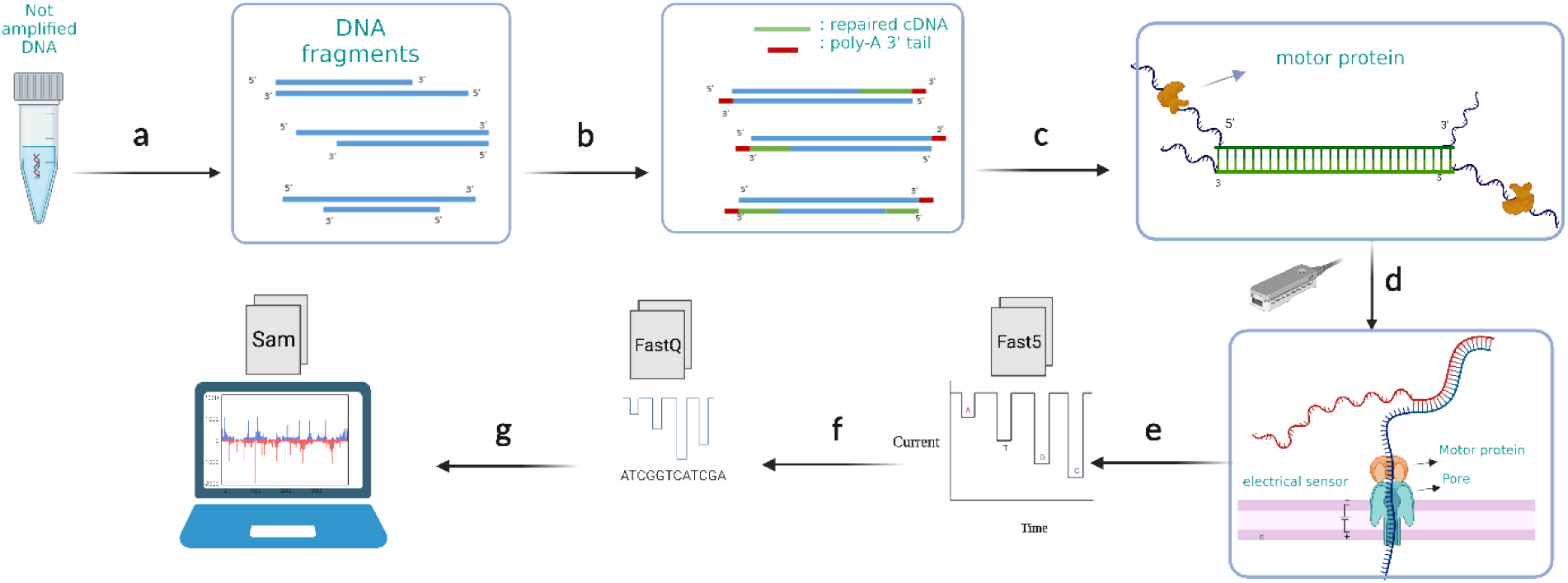
An overview of experimental workflow. a) DNA samples are amplified to higher concentrations using a panel of polymerases or plasmid DNA that is prepared from cells resulting in amplified products which have nicks and imperfections b) DNA fragment nicks are repaired, then end-prepped with poly-A tails at the 3’ ends. c) Adapters with motor proteins are ligated to the poly-A tailed end of the genes. d) The DNA library is loaded onto minION flow cells to start sequencing. Motor proteins attach to the pore proteins on the flow cell and push one strand of DNA through the chip pore creating a unique impedance signature for each base pair. e) Raw electrical signals are saved as Fast5 files. f) Raw signals are base called to assign oligonucleotides to each impedance event and these are converted to FastQ. g) Post sequencing analysis including alignment and enumeration of insertions, deletions, and substation of base pairs are performed on the nanopore base called data.

### Amplification methods

Following transformation of the pJL1-sfGFP plasmid into Top10 *E. coli*, colonies containing the plasmid were cultured and stored in a glycerol stock at −80°C. To initiate cell growth, a small amount of the stored cells was transferred to 10 ml of LB media and treated with Kanamycin to a final concentration of 50 μg/mL to selectively purify the cells with *pJL1-sfGFP*, due to the plasmid antibiotic resistance gene. The *E. coli* cells, containing the plasmid, were then cultured for 16 hours at 37°C with a rotation speed of 230 RPM. After the 16-hour incubation, the grown cells were collected, lysed, and the DNA was purified using the Promega Wizard Plus SV Miniprep DNA Purification System. The purified DNA was quantified using the Qubit dsDNA BR Assay, with the Qubit 4 Fluorometer, resulting in a typical yield of 250-330 ng/μL DNA concentration in 50 μL of nuclease-free water.

As described in detail in our prior works [13], all linear expression templates (DNA) were designed and optimized for expression in *E*.*coli* using IDT’s codon optimization and purchased from Integrated DNA Technologies (IDT) (Coralville, IA) as gBlock gene fragments. Sequences can be found in supporting information section 1. LET DNAs were suspended in nuclease free water to reach a final concentration of 10 ng/μl. All PCR amplifications including OneTaq polymerase (New England BioLabs, Ipswich, MA), Phusion High-Fidelity polymerase (New England BioLabs, Ipswich, MA), and Q5 High-Fidelity polymerase (New England BioLabs, Ipswich, MA) were done as stated in the manufacturer’s protocols. Resulting, amplified DNA was cleaned, purified, and concentrated using a Zymo purification kit (D4004) and the final concentration was calculated using a Qubit dsDNA BR Assay and with the Qubit 4 Fluorometer to reach at least 50 μl of DNA with an average of 70 ng/μl final concentration.

Rolling circle amplification (RCA) was done with the same LET as ordered above. Synthesized DNA was digested by HindIII-HF restriction enzyme (New England BioLabs, Ipswich, MA) by adding 5 μl of 10x buffer and 1 μl enzyme to 25 μl of purchased DNA with a concentration of 20 ng/μL; HindIII-HF was then inactivated by raising the solution temperature to 80°C for 20 min. T7 DNA ligase (New England BioLabs) ligated the sticky ends of the last step product to make circularized templates (CT). It was performed at 25°C for 1 h with the following reagents: 25 μl digested DNA from previous step, 25 μl of T7 DNA ligase 2x buffer, and 1 μL T7 DNA ligase enzyme. The circularized DNA templates were cleaned and purified using a Zymo purification kit and eluted in 50 μL of nuclease free water to get a yield of 30 ng/μL. Rolling circle amplification was done with a TempliPhi kit (GE Healthcare, Pittsburgh, PA) which works with 10 pg/μl concentration of CTs. Therefore, previous step samples were diluted 1000x and added with the kit’s protocols (1 μL template DNA, 5 μl sample buffer, 5 μl reaction buffer, 0.2 μl enzyme) and replicated in the 4 PCR tubes to get a higher amount of RCA amplified DNA. The reaction was done with overnight incubation time (12-14 h) and at 30°C. Nuclease free water (14 μL) was added to each PCR tube to dilute the viscous sample, mixed, and then purified with the Zymo kit and diluted in 50 μL of nuclease-free water to get to 70 ng/μL final concentration of DNA.

### Sequencing and DNA library preparation

100-200 fmol of amplified or not amplified DNA was prepared via the SQK-LSK114 or SQK-LSK110 sample preparation kit (ONT) to load on R10.4.1 or R9.4.1 flow cells. In short, DNAs are repaired with NEBNext FFPE module (New England BioLabs) and end-prepped with Ultra II End-prep module (New England BioLabs) at the same time. After purification and DNA quantity measurements (Qubit dsDNA BR Assay), adapter ligation and clean up was done following manufacturer’s protocols. Final DNA concentration was measured with the Qubit dsDNA BR Assay and 50 fmol of the DNA was loaded into the minION flow cells via following the ONT workflow instructions.

The principle of the device is to use the motor protein to push one strand of the DNA attached to it to protein pores and the flow cell’s sensor array. Each passing oligonucleotide causes a unique disruption of current which is the electrical signal output of the device. For each sample, around 20 – 40 Giga bases of data (2-6 million reads) were collected in the raw electrical format (Fast5) to be processed in the next steps.

### Illumina sequencing and heterogeneous samples preparation

Various mixtures of sfGFP LET and the folding reporter GFP (frGFP) LET were prepared by combining different ratios of DNA encoding for frGFP to sfGFP, specifically in proportions of 1:10, 1:100, and 1:1000. Subsequently, these samples underwent Illumina sequencing using the end-paired 150 bp MiSeq sequencer 300-cycle kit and Oxford Nanopore sequencing using R10.4.1 flow-cells and SQK-LSK114 kit. For the Illumina sequenced samples, quality control, coverage checking, and de novo genome assembly were carried out using QIAGEN CLC Genomics Workbench 23, applying default settings to ensure robust and consistent results.

### Base calling, mapping, and data analysis

Fast5 files from minION output, were base called to nucleotides in the format of fastQ files using the Guppy base caller,V 4.5.3 [20] on Ubuntu 20.04LTS (detailed codes and settings are in Supporting information section 7) with different Q-score filters and with 400bps and HAC settings for R10.4.1 and 450bps and HAC for R9.4.1. Alignment was done for both Illumina and Oxford Nanopore data with Minimap2 [21] using the preset parameters to align the fastQ and fasta files with the reference sequence. Minimap2 output (SAM format) was converted, sorted and indexed into BAM by Samtools program [22] and converted to CSV format by a custom code from our group (Supporting information section 7). The CIGAR string, which is a string that includes the data of matched, mismatched, inserted and deleted sequences and their position in the read sequence (example shown in Supporting information section 6) was analyzed to extract the error rates for each mutation by parsing the CIGAR string into the parts each operator in it is representing. The error rate for each mutation type (insertion, deletion, and substitution) was defined as the percentage of total mutations of each type normalized to the aligned length of the read data with the reference sequence (also equal to the error probability per base pair as a percentage). For the pJL1-sfGFP plasmid the maximum common sequence between the plasmid, and the linear template is solely considered as a reference sequence to map plasmid data with, which is from the T7 promoter to T7 terminator on pJL1-sfGFP.

### Error prone PCR

The reaction mixture is made of 50 μl nuclease free water, 10 μl 100 mM Tris-Cl, pH 8.3 (10 mM final), 2.5 μl 2 M KCl (50 mM final), 3.5 μl 200 mM MgCl2 (7 mM final), 3.5 μl 200 mM MgCl_2_ (7 mM final), 16 μl 25 mM dNTP mix (1 mM final), 2 μl 100 μM forward primer (2 μM final), 2 μl 100 μM backward primer (2 μM final), 10 μl 10 ng/μl template DNA (1 ng/μl final), 2 μl 25 mM MnCl_2_ (0.5 mM final), and 1 μl 5 U/μl OneTaq DNA polymerase (0.05 U/μl final). Each thermal cycle consists of three steps of a) 95°C for 1 min, b) 50°C for 1 min, (c) 72°C for 1 min. A 10 μl of reaction mixture is collected after 10 cycles and purified and cleaned with Zymo purification kit (D4004). The resulting DNA after 10 cycles got amplified with the OneTaq polymerase (New England BioLabs, Ipswich, MA) as the manufacturers protocol to reach a final concentration about 30-70 ng/μl measured by Qubit. The same procedure was continued after 15, 20, 25, and 30 cycles of error prone amplification with the reaction mixture.

### Cell free protein expression and emission spectra analysis

All cell-free reactions were conducted at 37°C in volumes of 15 μl using the following: 3.6 μl of 30% v/v *E. coli* extract (produced according to our previously published work on scalable extract [23], 5 μl of PANOx-SP system as supplement mixture (260 mM potassium glutamate, 8 mM magnesium glutamate, 4 mM potassium oxalate, 1.5 mM spermidine, 15 mM potassium phosphate (pH 7.0), 1.2 mM AMP, 0.86 mM each of GMP, UMP, and CMP, and 2 mM 19 amino acids with 1 mM tyrosine due to solubility) 5 nM of DNA template, and remaining volume with nuclease-free water to 15 μl total volume. Fluorescent protein sfGFP was expressed in a 384-well plate covered with a transparent film at 37°C for 10 h. The reactions were monitored using a SpectraMax iD3 microplate reader (Molecular Devices, USA) with a fluorescence reading every 30 min after the start of the reaction. sfGFP fluorescence was detected with an excitation/emission of 485/528 nm, a band-pass window of ±20 nm with auto gain. Emission peak data was gathered with a fixed excitation at 475 and an emission range from 495 nm to 600 nm, a band-pass window of ±20 nm, and auto gain. The resulting emission spectra was collected after 4 hours of reaction, the peak was detected and the fluorescence intensities after the peak were mirrored in order to make a symmetrical emission peak. Lorentizan function was fitted to the mirrored spectra and FWHM was extracted from the function (see the Supporting information section 7).

## Results

### Oxford Nanopore read quality and quantity

The genes were sequenced using Oxford Nanopore Technology sequencing flow cells and library preparation kits, specifically R10.4.1 paired with the SQK-LSK114 kit, and R9.4.1 paired with the SQK-LSK110 kit, for both amplified and non-amplified genes. The quantity of sequenced data varied depending on the library quality for both methods. It was previously reported that R10.4.1 generated half the amount of data compared to R9.4.1 in whole genome sequencing [24]. However, we observed that R10.4.1 produced more data in certain samples. For instance, in the case of the Phusion polymerase amplified sample, R9.4.1 generated 91% more sequencing data than R10.4.1. Conversely, when working with plasmids, R10.4.1 generated 193% more data than R9.4.1.

In terms of read accuracy, R10.4.1 outperformed R9.4.1 by yielding a greater amount of higher accuracy data. This is evident in the distribution of Q-scores (which can be converted to read accuracy using Equation 1) for read accuracy across various samples, as depicted in Supporting information Figure S1. When comparing the averages, each sample’s Q-score showed improvement ranging from 1.35 to 4.89. Notably, a significant observation was the ability of R10.4.1 to generate data with Q-scores exceeding 20 (Q20+), thus enabling a focus on high-quality data in subsequent analysis steps.

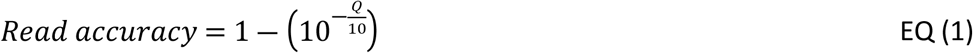

### Error rates in heterogeneous samples, Illumina vs Oxford Nanopore

Although short-read sequencing is recognized for its higher read accuracy compared to Oxford Nanopore, it has limitations in individual gene analysis. Short-read sequencing requires the assembly of longer sequences from numerous short DNA fragments. This process not only demands time and computational resources but may also lead to incomplete assembly in highly repetitive sequence regions[25]. Consequently, heterogeneity analysis is often conducted on short reads mapped to the reference genome. In this experiment, we aimed to explore the capability of NGS to detect heterogeneity within samples by mixing different ratios of the two genes, namely sfGFP and a modified folding reporter protein (frGFP) gene as the errored sequence which is designed to have has 3.741 % substitution, 1.011% deletions and 1.011% insertion error rates calculated based on the reference genes mapping compared to the sfGFP.

Initially, we conducted an analysis of *de novo* assembled sequences in an effort to generate longer contigs from MiSeq Illumina sequencing data, as shown in Supporting Table 1. Even in the case of the pure samples, which exhibited the longest contig at 734 base pairs (bps) for frGFP pure sample, it was insufficient to reach the full gene sequence length of 999 bps. Notably, for samples containing known, simulated impurities, the maximum contig length decreased. Additionally, the coverage of the sequenced DNA to the sfGFP reference genome was non-uniform, with observed lower coverage at the start and end regions of the gene. Through the analysis of error positions in frGFP and sfGFP sequences, highly erroneous areas were identified from the 774th to the 834th bp, coinciding with regions of low coverage. This phenomenon suppresses the average error rate by having reduced coverage at highly erroneous regions of the gene. (supporting information figure S-2)

The error rate analysis was conducted on both long-read and short-read sequences mapped to the sfGFP reference sequence, resulting in the calculation of average error rates for the mixtures, as shown in **Figure 2**. It’s worth noting that the vendor’s sfGFP gene, when mapped to the sfGFP DNA template sequence, exhibits a baseline error in both Illumina-sequenced and nanopore-sequenced data, indicative of previously reported inherent errors in the synthesized DNA from the vendor detected with the current read accuracy [25]. Another observation in Illumina is the abnormal error rates observed when frGFP was mapped to sfGFP for Illumina sequencing. These rates consistently reported smaller values for error rates across various error types. In contrast, the Oxford Nanopore sequencing method reported error rates for pure frGFP samples that were closer to the expected values (Fig 2b).

**Figure 2.**
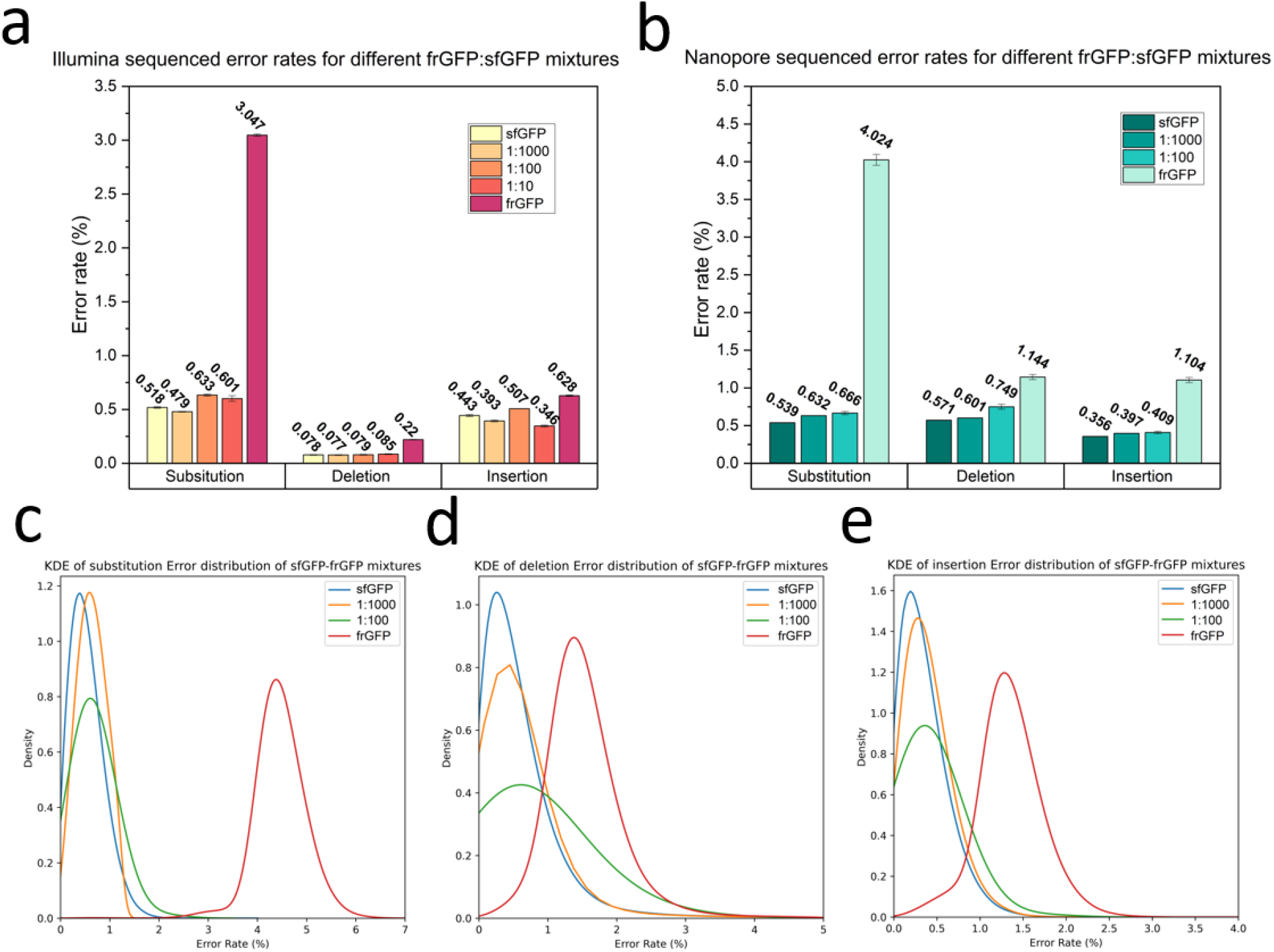
Benchmarking error rates observed in short-read vs. long-read sequencing approaches with known error-introduced heterogenous samples. Error rate averages for short-read Illumina sequenced data **(a)** and long-read Oxford Nanopore sequenced data **(b)** of different mixtures of pure sfGFP and errored frGFP sequences. Error bars show standard error and red solid lines show the expected percentage based on known ratios of pure and error templates. KDE fitted plots for the distribution of **c)** Substitution, **d)** Deletion, and **e)** Insertion errors for Oxford Nanopore sequenced samples with introduced error.

For the mixture analysis, the Illumina data displayed a convergence with the expected trend for deletions, while divergent patterns were observed in substitution and insertion assessments. By scrutinizing the distribution of error rates in the Illumina data, it became apparent that the majority of the errored sequences in the mixtures were sequences without any errors, with a small portion of sequences contributing to the errors in the averages (see Supporting information figure S2). Interestingly, even when Illumina sequenced pure frGFP sequences, the majority of reads exhibited no deletions or insertions errors, prompting an inquiry into the potential artificial error suppression within the Illumina sequencing pipeline during heterogeneity analysis which was not observed in the Oxford Nanopore sequenced data (supporting information Figure S4).

In contrast, Oxford Nanopore sequenced error rates across all sections matched expectations with the anticipated increasing trend as the ratio of frGFP elevated in the mixture (Fig 2b). Calculating the averages of errors in the samples and comparing them with the expected error rates for each mutation type, as determined from the values observed for pure sfGFP and frGFP for each sequencing method, revealed noteworthy differences. This analysis, detailed in the Supporting information tables S3 and S4, indicated that Illumina samples exhibited more pronounced divergence from the expected results in every section, further indicating the superior performance of long-read sequencing for heterogeneity analysis.

Kernel Density Estimation (KDE) was employed to analyze the distribution of errors of Oxford Nanopore sequencing, yielding valuable insights into each distribution. Specifically, KDE provided information about two key aspects: the peak value (mode of the fitted model) and the peak height, which signifies the sharpness of the peak and the homogeneity of the sample. As illustrated in **Figures 2c, 2d**, and **2e**, when the proportion of errored sequences in the mixture increased, the mode values for the error rates also increased, while the peak height decreased as the ratio of impurity increased. Notably, in the case of the pure frGFP sample, although the mode values exceeded those of the 1:100 mixture, indicating more error rates for the mode of the distribution, but the peak height indicated a higher level of homogeneity in the frGFP pure sample, further supporting the idea that peak height can be indicative of sample homogeneity.

### Error rate assessment using different read-quality filters

Oxford Nanopore data was collected with R9.4.1 and R10.4.1 flow cells to compare the effects of chip version and the read quality filter. First, we investigate the effect of different Q-score filters applied with the base caller to average error rates of different samples. It should be noted that only the linear templates could generate enough Q20 data for this analysis and therefore only non-amplified sfGFP and PCR products are analyzed in this section. RCA and plasmid samples had reduced read efficiency likely due to blocked pores caused by excess amounts of smaller DNA fragments. This would be resultant of digestion steps necessary to prepare these for Oxford Nanopore sequencing. By analyzing the read length of the sequenced data, smaller fragments were observed in the RCA and plasmid samples (Supplement figure S5) which would change the expected molar weight of each template and overshooting the correct molar window suggested by the vendor for sequencing (10-20 fmol of final prepared DNA library for the R10.4.1 flow cell). Three quality score filters were applied in this study: 1) the default setting on the Guppy base caller configured at a Q-score equal to 9 which filters out sequences with a read accuracy lower than 87.4%, 2) Q-score 15 (96.8% read accuracy) and 3) Q-score 20 (99% read accuracy). As depicted in **Figure 3a**, while increasing the Q-score filter resulted in decreased error rate values, the relative ranking between different polymerases remained consistent in substitution error assessment. However, for deletion (**Figure 3b**) and insertion errors (**Figure 3c**), the default Q-score setting exhibited significant variations in polymerase performance, with this variation diminishing as read accuracy increased. At Q-score 20, only OneTaq enzyme demonstrated statistically significant difference compared to non-amplified DNA, Phusion, and Q5. These findings emphasize the crucial role of read accuracy in benchmarking error rates. However, it’s important to note that increasing Q-score filters lead to a reduction in data volume. For example, by applying Q-score 15 filter, 20-50% of data passed the filter and by applying Q-score 20 between 0.6% - 0.01 % of data passed the required read quality (also see supporting figure S1 for Q-score distribution of samples). Therefore, a trade-off between read accuracy and data size is inevitable. Consequently, for further analysis of error rate distributions among all amplification samples, a Q-score of 15 was employed.

**Figure 3.**
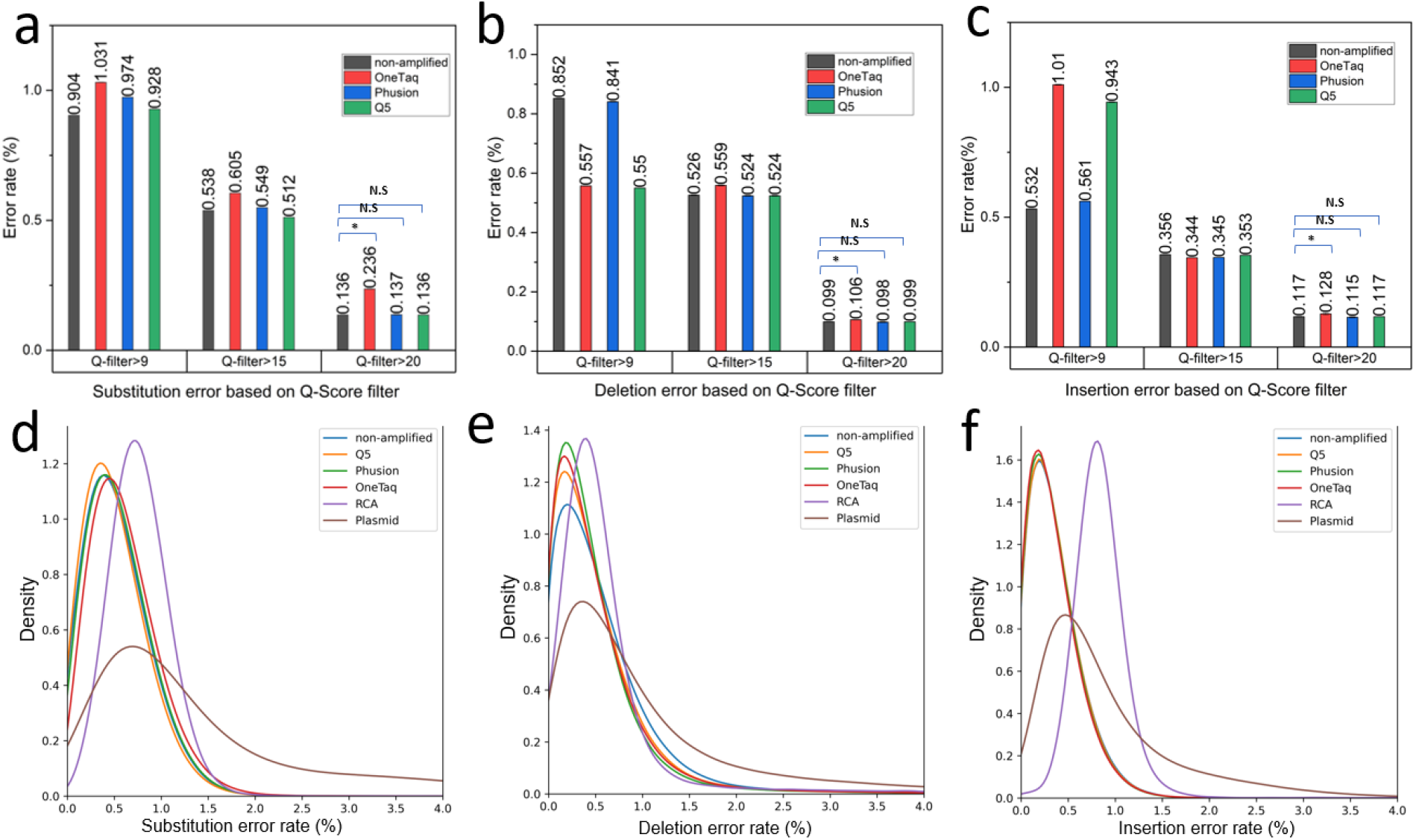
Comparing mutation error rate averages and distributions for different amplification methods. Average substitution **(a)**, deletion **(b)**, and insertion **(c)** error rates in samples amplified by various polymerases, filtered by fit accuracy (Q score). Distributions of substitution **(d)**, deletion **(e)**, and insertion **(f)** error rates between samples amplified by different methods.

### Amplified gene analysis

The panel of polymerases (OneTaq, Phusion, and Q5) were used to amplify the linear template gene using standard PCR as well as Phi-29 polymerase with RCA. pJL1-sfGFP plasmids were also amplified using a cell-based amplification, purified, and digested to make linear pJL1-sfGFP ready for sequencing. The results from the starting synthetic gene without amplification is also included in this section as a relative baseline for the errors assessed. Kernel density estimation models (KDE) were used to fit the distributions of each error type and visualize error rate profiles in the samples. For the substitution error rate (**Figure 3d**), the Q5 enzyme exhibits the lowest mode at 0.302%, while among the polymerases, it displays the highest peak. This observation suggests a more uniform error rate profile compared to the other polymerases.

Despite RCA having the highest peak across all samples, it also possesses the highest mode value of 0.723%. Consequently, it generates sequences with a homogeneous but elevated error rate. This could be due to errors introduced in the circularization steps. The plasmid substitution error distribution records the lowest peak, indicating greater heterogeneity in error rates among sequences produced by this amplification method compared to non-cell-based methods.

In the case of deletions (**Figure 3e)**, all polymerases demonstrate closely clustered error rate modes, ranging from 0.169% for OneTaq to 0.199% for Q5. The peak height of Phusion suggests its status as the most homogeneous polymerase in this panel. RCA, once again, exhibits a sharp peak but with a mode value of 0.38, while plasmid amplification results in the widest peak and the most heterogeneous sequences.

In insertions (**Figure 3f**), all polymerases exhibit close modes (0.200-0.203%) and peak heights. RCA stands out with the sharpest peak and the largest mode for error rate (0.816%). Plasmid amplification yields a mode value of 0.440, featuring the widest and lowest peak, once again showcasing the most heterogeneous amplification.

To assess the statistical significance of the error rate distributions, a bootstrapping[26] approach was employed for this analysis. This analysis tests the selection of multiple samples, each with a specified size, from two datasets. Following this, a Mann-Whitney U test was executed on each pair of bootstrapped samples. Over multiple iterations, the average p-value from the Mann-Whitney U test was computed, serving as the representative p-value for the comparison of the two distributions (see Supporting information section 5). All amplified samples exhibited statistical significance when compared to other samples, as determined by a 95% confidence level.

### Analysis of error location

Oxford Nanopore sequencing data readily allows for determining the locations of each type of mutation along the matched open reading frame (ORF) to determine if they are localized or randomly distributed. When plotted as stacked heatmaps (**Figure 4 a-c**) repeating patterns of mutation locations are apparent in the synthetic (not-amplified) DNA, as well as the OneTaq, Q5, RCA, and Phusion amplified DNA. This indicates that the mutation findings were not random errors from the Oxford Nanopore sequencer but rather conserved areas of high error rates in the original synthesized DNA. The plasmid, which is amplified via cellular growth, has a pattern of errors distinct from the linear templates. Only the plasmid portion common to the LET sequences, namely between the T7 promoter and terminator or from 185^th^ bp to 1756^th^ bp backward on 3’, were analyzed; this results in 915 bps of the total pJL1-sfGFP being analyzed and plotted. More isolated, distinct bands in the plasmid insertion and deletion mutation heatmaps are observed, which indicates that these plasmid mutations are isolated to a few, specific hotspots. The location of the insertion and deletion hotspots can be projected to the pJL1-sfGFP plasmid map (Supporting information section 1) and are determined to be before the final beta-barrel of the protein (GFP-11, Addgene Sequencing Result #199339) and after the sfGFP protein gene, respectively. It is not evident what the mechanism is for these localized errors, but it is clear Oxford Nanopore sequencing can identify them and help screen DNA for potential errors prior to downstream uses such as expression templates in CFE.

**Figure 4.**
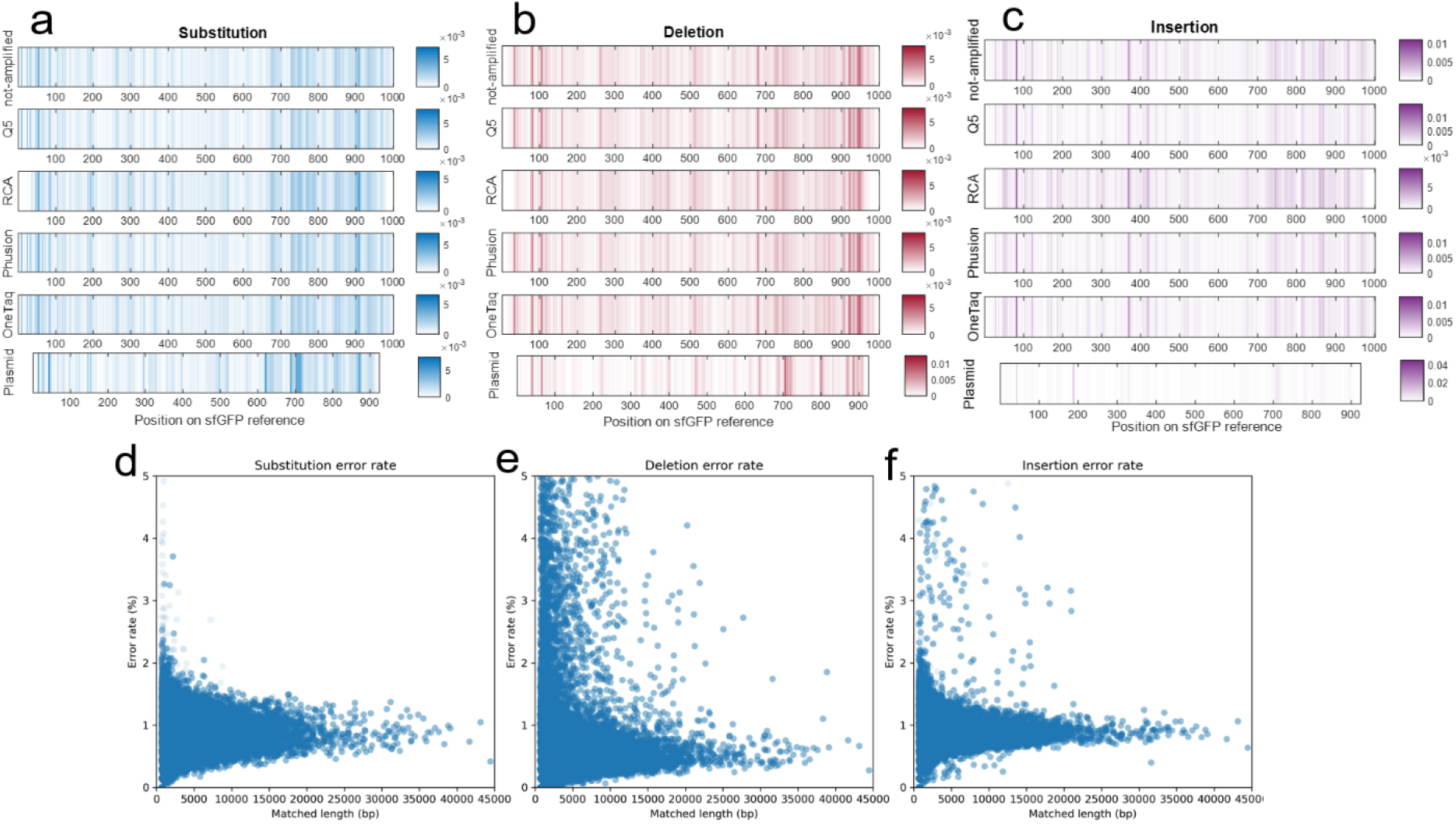
Location of mutations within the template and dependency of error rates to RCA product length: **a)** substitution, **b)** deletion, and **c)** insertion on the sfGFP reference gene. The intensity of colors corresponds to the likelihood of mutation occurrence (the color axis represents the probability of mutation at each base pair or the count of observed mutations at that location, relative to the total number of read base pairs at that location), with darker shades indicating higher probability spots for mutations. **d)** Substitution **e)** deletion **f)** and insertion error rate distributions of RCA products based on the length of the matched sequence length.

Also, by mapping the RCA sequences onto a repeated reference sfGFP sequence, error rates normalized by the overall matched length of the sequence can be plotted to analyze the dependency of error rates observed in RCA samples to the length of RCA product. As depicted in **Figure 4e-f**, the error rate profiles for longer amplified DNAs closely follow the mode of errors observed in shorter fragments observed in the distribution plots (**Figure 3d-f**). Moreover, the distribution is observed to be linear with respect to length which suggests that the Phi29 polymerase contributes little to the observed error rate, and rather duplicates errors found in the starting, circularized template (in our case, caused by initial PCR of the starting template). If the Phi29 enzyme contributed significantly to the error, it would cause more error as the template increased in length which would cause this observed peak to slant upwards in Fig 4d-f. Therefore, RCA can be a valid CFE template generation technique if care is given to the initial quality of starting sequence (using a high-quality polymerase for initial amplification).

### Secondary evidence of mutation heterogeneity from cell-free expressed sfGFP spectrums

To corroborate the sequencing results and link to CFE performance, sfGFP was expressed in cell free extracts using the panel of amplified DNA linear templates, plasmid, and synthetic DNA from vendor (no amplification). We hypothesized that the emission spectrum data, from 495 nm to 600 nm, would broaden as more mutations were made to the protein, caused by imperfections in the beta barrel structure and thus would vary the photophysical properties of this standard reporter protein. Different mutants of sfGFP have shown varied emission and excitation maximum wavelengths. For example two amino acid mutations (N149Y/Q204H) on sfGFP can result in producing pH-tdGFP which has an emission peak (Em λ) of 515 nm that is red shifted 5 nm in comparison with 510 nm Em λ of sfGFP [27]. As an example of blue shifting of Em λ, usGFP with two mutations on sfGFP original sequence (Q69L/N164Y) has shown an Em λ of 508 nm [28].Even one mutation can change sfGFP to a yellow sfYFP (T203Y), a cyan sfCFP (Y66W) or a blue sfBFP (Y66H) [29]. The intensity of the spectrum (overall fluorescence count) would not be a good measure, as this can vary between LET and plasmid simply due to the efficiency of transcription and competing degradation of linear templates.

Excitation wavelength was held constant and CFE replicates were done in 8-10 wells for each amplification method due to inherent variability of this technique [30]. The widths of the emission peaks (around 510 nm) were determined using the full width at half maximum (FWHM) of the fitted Lorentzian function to the mirrored data (Fig 5a). The reason for mirroring data after the peak is 1) emission data near to the excitation cannot be collected and 2) to construct a symmetrical peak suitable for fitting symmetric models such as Lorentzian and Gaussian. As proof of concept, sfGFP genes were first subjected to an error-prone PCR step with varying cycle numbers and then normally amplified to reach a suitable concentration for performing CFE. As shown in Figure 5b, with an increase in the cycles of error-prone PCR, the FWHM average of the samples also increases, indicating a broadening effect on the emission spectra peak. Analyzing FWHM averages for different amplification methods (Figure 5c), it is observed that RCA and OneTaq-amplified genes exhibited the widest peaks. Non-amplified genes, as well as those amplified with Q5, Phusion, and the plasmid, showed the narrowest peaks, and they were not statistically different from each other. This suggests the production of more homogenous sfGFP using Q5 and Phusion for linear amplifications, as well as using the plasmid template. Additionally, it’s important to note that non-amplified genes also exhibited small variations in FWHM values (Fig 5 b,c). This variability can be accounted for by variability in measurement technique (data taken on different days) and some variability caused by using different batches of fresh DNA from the vendor.

**Figure 5.**
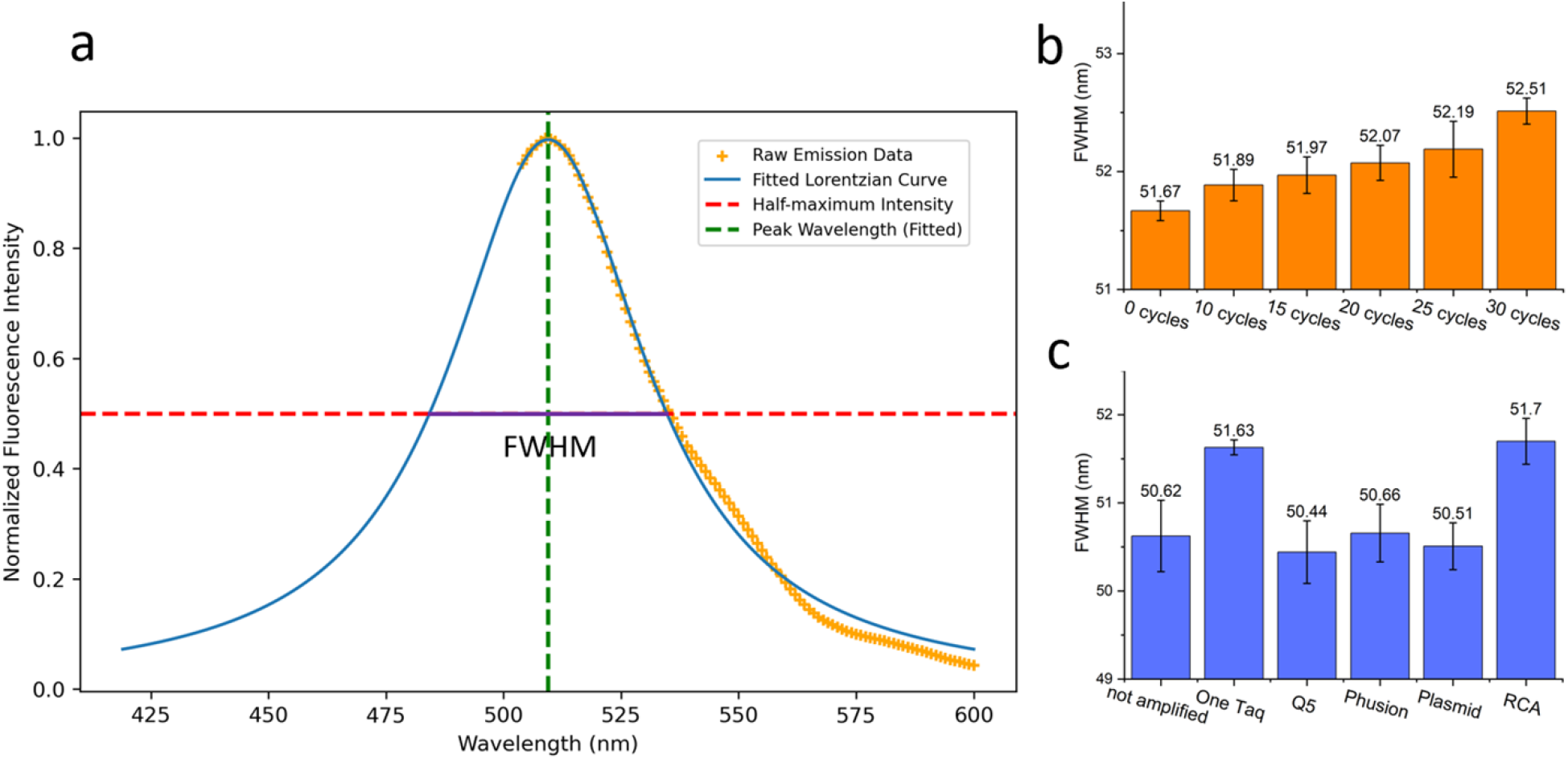
Emission spectra analysis. **a)** Depiction of emission peak half width measurement using mirrored emission spectral data fitted by a Lorentzian function. The green dotted line is the peak wavelength and the red dotted line is the half-maximum intensity. The purple line indicated the FWHM for the spectra. **b**) A comparison between the emission spectra peak’s FWHM of different sfGFP genes with 0,10, 15, 20, 25, 30 cycles of error prone-PCR. Each bar is the average and the error bars are the standard error of the samples. **c)** sfGFP emission spectra peak’s FWHM of different amplification methods.

## Discussion

We first investigated the efficacy of NGS compared to Oxford Nanopore sequencing for heterogeneity analysis. Our results revealed that NGS data obtained by MiSeq Illumina, specifically with 150 bps end-paired reads, is not able to achieve heterogeneity assessment of our intentional error-induced samples. Several potential explanations exist for these observations. First, the analysis of errors in NGS data is constrained by short-read mapping, limiting the ability to filter smaller fragments and impurities from the synthesized DNA by implementing a minimum length requirement for the parent gene. Another possible explanation could be the uneven coverage of sequenced reads across the reference genome. Areas with higher error rates tend to have lower coverage, which might lead to a reduction in average error rates due to a bias toward reading more low-error regions. Additionally, previous research has indicated that Illumina sequencing read accuracy can vary due to factors such as library preparation and enrichment PCR [31]. Also in end-paired NGS data, a higher error rate in the second read (R2) has been observed, prompting questions about the overall accuracy of NGS for error heterogeneity assessments [32].

Another important observation was the influence of sequencing errors as a potential source of bias in evaluating DNA quality specially in the ONT samples. Initial results obtained from R9.4.1 flow cells demonstrated biased outcomes in our error analyses. However, by adopting a more precise flow cell and chemistry kit, specifically R10.4.1 flow-cell and SQK-LSK114 kit, we were able to investigate the impact of read accuracy on error rate assessments. This led to the identification of read accuracy filtering as a crucial requirement for ensuring unbiased error assessments [33]. While the results obtained with an applied Q-score of 15 (read accuracy 96.8%) were in line with those at Q-score 20 (accuracy 99%) for the analysis of various polymerases’ fidelity, the requirement for higher-quality sequencing data becomes essential to conclusively validate results, especially when dealing with polymerases exhibiting small differences in fidelity. The need for high-volume data coupled with superior read accuracy highlights the importance of leveraging advanced sequencing technologies like PacBio for a comprehensive confirmation of these findings.

A recent study found no significant difference in errors introduced to the synthesized gene by Q5 and Phusion polymerases, as measured using NGS. The source of errors for the sequences was identified as the synthesized gene itself rather than the amplified sequences [25]. This discovery aligns with our observation that there is no statistical significance in the average error rates between non-amplified DNA, Phusion, and Q5 amplified DNAs when a Q-score 20 filter was applied in our Oxford Nanopore pipeline, elevating the read accuracy to 99%. The feasibility of obtaining such findings from Oxford Nanopore data was not realized until recent advances improved the read quality of this sequencing method. Among the PCR amplification methods, only OneTaq enzyme showed higher error rates in averages when a Q-score 20 filter was applied, which is in line with literature comparing this widely used enzyme to newer, high fidelity enzymes such as Phusion and Q5 [12, 34, 35].

Another analysis that can only be done with long-read sequencing, is the analysis of heterogeneity of errors between individual sequences of DNA read by nanopores. It enables getting useful information about the distribution of errors between the sample, rather than calculating an average of errors, calculated by typical NGS approaches. This analysis is specifically useful when analyzing repeated sequences like RCA products, to analyze errors in each long-repeated sequences individually. In our test result, we observed that the longer RCA sequences have the same error rates as the shorter ones, confirming Phi29 high fidelity for the long amplifications.

Regarding RCA, both R9.4.1 and R10.4.1 flow-cells resulted in a relatively low yield of sequencing data which is attributed to the broad spectrum of fragment lengths observed in the read length distribution after sequencing. This diversity posed a challenge to accurately determine the molar basis amount of DNA required to optimize library preparation, such as adapter ligation. Additionally, the abundance of smaller fragments could potentially obstruct the nanopores, contributing to the observed lower sequencing yield in RCA sequencing. Also, the emission peak sharpness analysis reported a wide peak for RCA which confirms the Oxford Nanopore reported error rates for the RCA. This error location pattern suggests non-random errors in the amplification process, pointing to the probability of the initial circular template as the main source of errors rather than the Phi29 DNA polymerase itself. This conclusion is also supported by analyzing the error location pattern being similar to the non-amplified gene. The increased error may be introduced during circular template assembly, including the digestion of synthesized DNA and the ligation of nicked fragments to build the circular template. Fragments of incompletely synthesized DNA may contribute in this step, forming errored circular templates. Therefore, pre-amplifying with high-fidelity enzymes, such as Q5, as opposed to OneTaq [6], is considered the optimal strategy for RCA compared to directly making circular from the vendor DNA [36].

By analyzing the error locations on the pJL1-sfGFP Plasmid, it is shown that they are focused in a few specific locations for insertions and deletions (hotspots). Only the insertion error hotspot around 200 is inside the sfGFP coding region and the bands around 700 for deletions and insertions are outside of the protein coding region of plasmid and would not affect protein functionality, which we see evidence of in that the Oxford Nanopore data shows increase in error for the plasmid but not in the fluorescent data in the CFE experiment (tight emission curves). Regarding the origin of elevated errors in plasmids, a plausible explanation could be the observed increase in error rates in bacterial genomic and plasmid DNA in the presence of antimicrobial threats. This phenomenon may enable the organism to undergo mutations, potentially leading to the development of antimicrobial resistance genes [37]. Another explanation is that mismatch repair is greatly governed by the number of the errors in the plasmid; this repair mechanism decreases with an increase in other mismatches present in the DNA, especially deletions and insertions that result in marginal or even the lack of the mismatch repair [38]. The initial assembly of the plasmid using linear templates would affect this quality. Also, since the plasmid DNA is prone to methylation, a methylation-aware mode of Oxford Nanopore base calling could improve the quality of the sequence[39], however such analysis is not year readily available and beyond the scope of this work.

## Conclusion

In conclusion, this study highlights the potential of Oxford Nanopore sequencing as a useful tool for assessing amplified DNA quality especially when PacBio is beyond budget or geographic access; DNA homogeneity is a critical parameter in biotechnological laboratories globally, especially for synthetic biologists that require a high-quality genetic blueprint for their engineered protein product or circuit. Through error profile analysis, the high-fidelity polymerases Phusion and Q5 are found to perform best, offering robust amplification of linear templates for applications such as cell free protein expression. RCA with Phi29 polymerase is also a viable technique if the starting circular template has low error. Oxford Nanopore sequencing data also readily allows for assessment of mutation location. In the case of the sfGFP plasmid, the majority of deletions and insertions were found to occur downstream of the GFP coding sequence, thus not affecting the fluorescence properties. Alternatively, we demonstrated how sfGFP heterogeneity could be measured using emission peak width measurement. Oxford Nanopore can be used to measure performance of new template synthesis and amplification techniques as they are discovered and provide a means for benchmarking quality prior to CFE production.

## References

1. Silverman, A.D., A.S. Karim, and M.C. Jewett, Cell-free gene expression: an expanded repertoire of applications. Nature Reviews Genetics, 2020. 21(3): p. 151–170.

2. Swartz, J.R., Transforming biochemical engineering with cell-free biology. AIChE Journal, 2012. 58(1): p. 5–13.

3. Karig, D.K., Cell-free synthetic biology for environmental sensing and remediation. Current opinion in biotechnology, 2017. 45: p. 69–75.

4. Dopp, J.L., D.D. Tamiev, and N.F. Reuel, Cell-free supplement mixtures: Elucidating the history and biochemical utility of additives used to support in vitro protein synthesis in E. coli extract. Biotechnology advances, 2019. 37(1): p. 246–258.

5. McSweeney, M.A. and M.P. Styczynski, Effective use of linear DNA in cell-free expression systems. Frontiers in Bioengineering and Biotechnology, 2021. 9: p. 715328.

6. Dopp, J.L., et al., Rapid prototyping of proteins: Mail order gene fragments to assayable proteins within 24 hours. Biotechnology and Bioengineering, 2019. 116(3): p. 667–676.

7. Shevelev, I.V. and U. Hübscher, The 3′–5′ exonucleases. Nature reviews Molecular cell biology, 2002. 3(5): p. 364–376.

8. Bębenek, A. and I. Ziuzia-Graczyk, Fidelity of DNA replication—a matter of proofreading. Current genetics, 2018. 64(5): p. 985–996.

9. Reha-Krantz, L.J., DNA polymerase proofreading: Multiple roles maintain genome stability. Biochimica et Biophysica Acta (BBA)-Proteins and Proteomics, 2010. 1804(5): p. 1049–1063.

10. Ricardo, P.C., E. Françoso, and M.C. Arias, Fidelity of DNA polymerases in the detection of intraindividual variation of mitochondrial DNA. Mitochondrial DNA Part B, 2020. 5(1): p. 108–112.

11. Cline, J., J.C. Braman, and H.H. Hogrefe, PCR fidelity of pfu DNA polymerase and other thermostable DNA polymerases. Nucleic acids research, 1996. 24(18): p. 3546–3551.

12. Potapov, V. and J.L. Ong, Examining sources of error in PCR by single-molecule sequencing. PloS one, 2017. 12(1): p. e0169774.

13. Dopp, J.L. and N.F. Reuel, Rapid, Enzymatic Methods for Amplification of Minimal, Linear Templates for Protein Prototyping using Cell-Free Systems. JoVE (Journal of Visualized Experiments), 2021(172): p. e62728.

14. Esteban, J.A., M. Salas, and L. Blanco, Fidelity of phi 29 DNA polymerase. Comparison between protein-primed initiation and DNA polymerization. Journal of Biological Chemistry, 1993. 268(4): p. 2719–2726.

15. Shendure, J. and H. Ji, Next-generation DNA sequencing. Nature biotechnology, 2008. 26(10): p. 1135–1145.

16. Lin, B., J. Hui, and H. Mao, Nanopore technology and its applications in gene sequencing. Biosensors, 2021. 11(7): p. 214.

17. Tyler, A.D., et al., Evaluation of Oxford Nanopore’s MinION sequencing device for microbial whole genome sequencing applications. Scientific reports, 2018. 8(1): p. 1–12.

18. Wang, Y., et al., Nanopore sequencing technology, bioinformatics and applications. Nature biotechnology, 2021. 39(11): p. 1348–1365.

19. Luo, J., et al., Systematic benchmarking of nanopore Q20+ kit in SARS-CoV-2 whole genome sequencing. Frontiers in microbiology, 2022: p. 4059.

20. Wick, R.R., L.M. Judd, and K.E. Holt, Performance of neural network basecalling tools for Oxford Nanopore sequencing. Genome biology, 2019. 20(1): p. 1–10.

21. Li, H., New strategies to improve minimap2 alignment accuracy. Bioinformatics, 2021. 37(23): p. 4572–4574.

22. Danecek, P., et al., Twelve years of SAMtools and BCFtools. GigaScience, 2021. 10(2).

23. Dopp, J.L. and N.F. Reuel, Process optimization for scalable E. coli extract preparation for cell-free protein synthesis. Biochemical Engineering Journal, 2018. 138: p. 21–28.

24. Ni, Y., et al., Benchmarking of Nanopore R10. 4 and R9. 4.1 flow cells in single-cell whole-genome amplification and whole-genome shotgun sequencing. Computational and Structural Biotechnology Journal, 2023. 21: p. 2352–2364.

25. Masaki, Y., Y. Onishi, and K. Seio, Quantification of synthetic errors during chemical synthesis of DNA and its suppression by non-canonical nucleosides. Scientific Reports, 2022. 12(1): p. 1–9.

26. Berrar, D., Introduction to the Non-Parametric Bootstrap. 2019.

27. Roberts, T.M., et al., Identification and Characterisation of a pH-stable GFP. Scientific reports, 2016. 6(1): p. 1–9.

28. Yong, K. and D. Scott, Rapid directed evolution of stabilized proteins with cellular high-throughput encapsulation solubilization and screening (CHESS). Biotechnology and Bioengineering, 2015. 112(3): p. 438–446.

29. Pédelacq, J.-D., et al., Engineering and characterization of a superfolder green fluorescent protein. Nature biotechnology, 2006. 24(1): p. 79–88.

30. Dopp, J.L., Methods to reduce variability in E. Coli-based cell-free protein expression experiments,. 2019. Volume 4: p. 204–211.

31. Ma, X., et al., Analysis of error profiles in deep next-generation sequencing data. Genome biology, 2019. 20: p. 1–15.

32. Tan, G., et al., Long fragments achieve lower base quality in Illumina paired-end sequencing. Scientific reports, 2019. 9(1): p. 2856.

33. Sanderson, N., et al., Comparison of R9. 4.1/Kit10 and R10/Kit12 Oxford Nanopore flowcells and chemistries in bacterial genome reconstruction. BioRxiv, 2022: p. 2022.04. 29.490057.

34. Filges, S., et al., Impact of polymerase fidelity on background error rates in next-generation sequencing with unique molecular identifiers/barcodes. Scientific reports, 2019. 9(1): p. 3503.

35. Lee, D.F., et al., Mapping DNA polymerase errors by single-molecule sequencing. Nucleic acids research, 2016. 44(13): p. e118–e118.

36. Hadi, T., et al., Rolling circle amplification of synthetic DNA accelerates biocatalytic determination of enzyme activity relative to conventional methods. Scientific Reports, 2020. 10(1): p. 10279.

37. Meyerovich, M., G. Mamou, and S. Ben-Yehuda, Visualizing high error levels during gene expression in living bacterial cells. Proceedings of the National Academy of Sciences, 2010. 107(25): p. 11543–11548.

38. Marinus, M., DNA mismatch repair. EcoSal Plus, 2012. 5(1): p. 10.1128/ecosalplus. 7.2. 5.

39. Yuen, Z.W.-S., et al., Systematic benchmarking of tools for CpG methylation detection from nanopore sequencing. Nature communications, 2021. 12(1): p. 3438.

